# Hopanoids confer robustness to physicochemical variability in the niche of the plant symbiont *Bradyrhizobium diazoefficiens*

**DOI:** 10.1101/2021.08.31.458470

**Authors:** Elise Tookmanian, Lisa Junghans, Gargi Kulkarni, Raphael Ledermann, James Saenz, Dianne K. Newman

## Abstract

Climate change poses a threat to soil health and agriculture, but the potential effects of climate change on soil bacteria that can help maintain soil health are understudied. Rhizobia are a group of bacteria that increase soil nitrogen content through a symbiosis with legume plants. The soil and symbiosis are potentially stressful environments, and the soil will likely become even more stressful as the climate changes. Many rhizobia within the bradyrhizobia clade, like *Bradyrhizobium diazoefficiens*, possess the genetic capacity to synthesize hopanoids, steroid-like lipids similar in structure and function to cholesterol. Hopanoids are known to protect against stresses relevant to the niche of *B. diazoefficiens*. Paradoxically, mutants unable to synthesize the extended class of hopanoids participate in similarly successful symbioses compared to the wild type, despite being delayed in root nodule initiation. Here, we show that in *B. diazoefficiens*, the *in vitro* growth defects of extended hopanoid deficient mutants can be at least partially compensated for by the physicochemical environment, specifically by optimal osmotic and divalent cation concentrations. Through biophysical measurements, we show that extended hopanoids confer robustness to environmental variability. These results help explain the discrepancy between previous *in vitro* and *in planta* results and indicate that hopanoids may provide a greater fitness advantage to rhizobia in the variable soil environment than the more controlled environment within root nodules. To improve the legume-rhizobia symbiosis through either bioengineering or strain selection, it will be important to consider the full lifecycle of rhizobia, from the soil to the symbiosis.

**Importance:** Rhizobia, such as *B. diazoefficiens*, play an important role in the nitrogen cycle by making nitrogen gas bioavailable through symbiosis with legume plants. As climate change threatens soil health, this symbiosis has reentered the spotlight as a more sustainable source of soil nitrogen than the energy-intensive Haber-Bosch process. Efforts to use rhizobia as biofertilizers have been effective; however, long term integration of rhizobia into the soil community has been less successful. This work represents a small step towards improving the legume-rhizobia symbiosis by identifying a cellular component—hopanoid lipids—that confers robustness to environmental stresses rhizobia are likely to encounter in soil microenvironments as sporadic desiccation and flooding events become more common.

## Introduction

The soil is a precious ecosystem. The health of the soil – measured by organic matter and nutrient content, moisture retention, and the microbial community – predicts how well plants will grow (1). While practices to maintain healthy soil have been known for centuries, as agriculture faces the threat of climate change, these sustainable land management practices have garnered new attention as potential climate change mitigation strategies (2, 3). Crop rotation is an ancient land management strategy that restores nutrients to the soil. Legumes, such as soybean, peanut, and alfalfa, are an important component of crop rotations because they increase soil nitrogen content, reducing reliance on synthesized nitrogen fertilizers (4). However, legumes cannot do this alone: they rely on a symbiosis with a group of polyphyletic soil bacteria called rhizobia (5, 6). This symbiosis is a very close interaction, with the bacteria living intracellularly in specialized *de novo* organs called root nodules. Low pH, low oxygen, and elevated osmolarity are maintained within the nodule environment (7–9). While in some ways stressful for the bacteria, this environment favors bacterial conversion of nitrogen gas to bioavailable ammonia which eventually is exchanged for reduced carbon in the form of dicarboxylic acids.

One way to improve legume use in crop rotations as a sustainable nitrogen fertilizer is to improve the efficiency of the legume-rhizobia symbiosis. Rhizobia strains with greater symbiotic efficiency, as measured by nitrogen fixation rates and legume growth, have been isolated and applied to the soil or to legume seeds. This strategy has been used successfully at scale with legume crops such as soybean in Brazil (10–12). However, the rhizobia often fail to stably integrate into the soil community, so these inoculations must be repeated each year (13–16). Beyond the symbiosis itself, the rhizobia can have positive effects on the plant when living in the soil, such as relieving salinity stress and increasing water and nutrient uptake (17–19). These positive effects will become even more important as the climate changes, leading to drastic changes in precipitation and thus soil water potential and osmotic strength. To successfully use legumes in crop rotations to increase soil nitrogen content as the climate changes, rhizobia must be successful in both the symbiosis and surviving the soil environment. The fact that these bacteria are ubiquitous indicates that they have strategies to accomplish both goals (20).

As recently discussed, possession of an adaptable outer membrane may provide an important fitness advantage in this context (9). Rhizobia produce modified lipid A, a major component of the lipopolysaccharides (LPS) that make up the outer leaflet of the outer membrane (21, 22). These modifications trend toward increasing the hydrophobicity and other cohesive interactions that lead to a more robust outer membrane (23, 24). Additionally, a subset of rhizobia, mostly within the *Bradyrhizobia* clade, make hopanoids, a class of sterol-like lipids, which maintain resistance to environmental stressors such as pH, temperature, antibiotics and foster successful symbioses (25, 26). Some *Bradyrhizobia* also attach hopanoids to lipid A, which appears to act as a hydrophobic hook into the inner leaflet of the outer membrane (27–29). This <u>ho</u>panoid attached to <u>l</u>ipid <u>A</u> (HoLA) is synthesized from extended hopanoids, a subclass of hopanoids with an added hydrophilic tail (Figure 1) (25).

**Figure 1.**
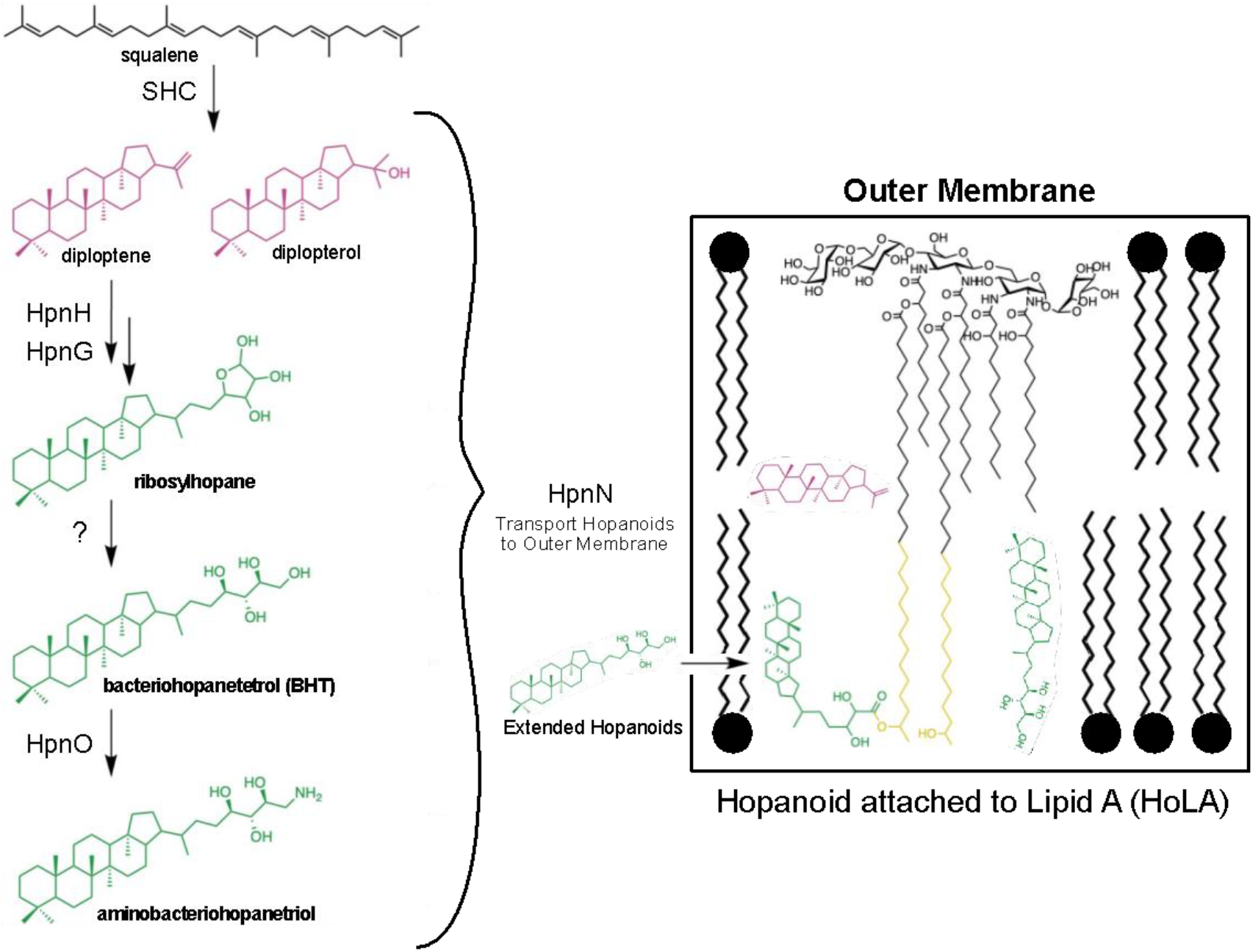
Hopanoid biosynthesis and HoLA structure. The biosynthesis of hopanoids is shown on the left from squalene to the unextended hopanoids (magenta) to the extended hopanoids (green). All of these hopanoids may be methylated at the C-2 position by the hopanoid methylase HpnP. These hopanoids are transferred to the outer membrane by the HpnN hopanoid transporter. Extended hopanoids are attached to the very long chain fatty acid (yellow) on lipid A to create HoLA which also resides in the outer membrane.

Intriguingly, generating a mutant that is unable to make any type of hopanoid by removing the first committed step in hopanoid biosynthsis (Δ*shc*) has evaded realization in *Bradyrhizobium diazoefficiens*, pointing to an essential role for hopanoids in this strain (25). Removing the ability to synthesize C_35_ or “extended” hopanoids (Δ*hpnH*), however, was achieved, and has a large effect on the fitness of *B. diazoefficiens in vitro* (25, 30). The Δ*hpnH* mutant manifests growth defects at high osmolarity and is unable to grow under low pH or microaerobic conditions – all conditions thought to characterize the root nodule. While the Δ*hpnH* mutant exhibits defects *in planta*, especially in root nodule initiation, the nitrogen fixation rate in symbiosis with the tropical legume *Aeschynomene afraspera* when normalized for nodule dry weight is not significantly different between the Δ*hpnH* mutant and WT and the majority of Δ*hpnH*-infected nodules grow at rates comparable to the WT (30). Given the growth defects of the Δ*hpnH* mutant observed *in vitro* in the presence of environmental stresses expected within the root nodule, these results were surprising. Here, we investigate this paradox by exploring the nuanced interplay between the lack of extended hopanoids and an environmentally relevant concentration range of osmolytes and cations.

## Materials and Methods

### Bacterial strains, Culture Media, and Chemicals

All strains used in this study are described in Table S1 in the supplemental material. All strains were grown aerobically with shaking at 250 rpm. *Escherichia coli* strains were grown in lysogeny broth (LB) at 37°C (31). *B. diazoefficiens* strains were grown at 30°C in either peptone-salts-yeast extract medium with 0.1% arabinose (PSY) (32, 33) or arabinose-gluconate medium (AG) (34, 35). The pH of the AG medium was adjusted to pH 6.6 using a NaOH solution or to pH 5 using a HCl solution. For pH 5 AG media, the HEPES buffer component was replaced with 5.5 mM MES buffer for a total of 11 mM MES. A low divalent cations pH 5 AG media was made containing 45 μM CaCl_2_ and 400 μM MgSO_4_. Inositol, sorbitol, CaCl_2_, or MgCl_2_ were added to the appropriate base medium. For induction of the cumate inducible promoter, cumate was added to liquid or solid medium for a final concentration of 25 μM from a 400x stock solution in ethanol (36–39). Agar plates were made containing 1.5% (w/v) agar. Antibiotics were used for selection at these concentrations (μg/ml): spectinomycin (Sp), 100; kanamycin (Km), 100; tetracycline (Tc), 20 for liquid cultures of *B. diazoefficiens* and 50 for plates for *B. diazoefficiens* and for *E. coli*. Chemicals were purchased from Sigma-Aldrich unless otherwise noted: glycerol (VWR), Hepes (Gold BioTechnology), sodium chloride yeast extract, magnesium sulfate, magnesium chloride, sorbitol, sodium hydroxide (Fisher Scientific), arabinose (Chem-Impex International, Inc), sodium sulfate (Mallinckrodt Chemical), peptone, and agar (Becton Dickinson).

### DNA methods, plasmid construction, and transformation

All plasmid constructions and primers used in this study are described in Table S1 in the supplemental material. Standard methods were used for plasmid DNA isolation and manipulation in *E. coli* (40). The strong constitutive promoter P*_rrn-mut2_* (41) was annealed from oligonucleotides Prrn-mut2_f and Prrn-mut2_r and cloned into *Hpa*I/*Bsr*GI-digested pQH2, resulting in plasmid pQH2-Prrn-mut2. The resulting inducible system (containing P*_bla-mut1T_-cymR\** and P*_rrn-mut2_* flanked by by *cuO*) was subsequently excised with *Spe*I and *Pci*I and ligated into pRJPaph-lacZYA prepared with *Spe*I and *Nco*I, resulting in plasmid pRJPcu1-lacZYA. The PCR product of the *shc* gene was cloned into the pRJPcu plasmids respectively to obtain expression plasmids. The pRJPcu-*shc* plasmid was mobilized into WT, followed by the pGK302, the markerless deletion vector to delete *shc* (blr3004) (42). Plasmids were mobilized by conjugation from *E. coli* S17-1 into *B. diazoefficiens* strains as previously described with the following modifications (43). The pRJPcu-*shc* plasmid was stably integrated as a single copy into the *scoI* downstream region of *B. diazoefficiens* as described previously (44).

### Induction conditions and reporter activity measurements

Cultures were grown in PSY to a mid-exponential phase and induced with 25 μM (final concentration) cumate or pure ethanol (the solvent for cumate) for controls. For quantitative LacZ assays, cells were centrifuged (5000 x g) and washed twice. β-Galactosidase assay was done as previously described (31). One biological replicate was defined as an independent culture, each replicate was assayed in technical duplicates of which the arithmetic mean was used for final data plotting.

### Streaking strains

Liquid cultures were grown from plates to early stationary phase (OD 0.9-1.2 by Beckman Coulter UV-VIS) in AG media. The cultures were spun down and resuspended to OD_600_ 0.5 in fresh media. 10 μL of culture was spotted on each plate and spread using a sterilized spreader.

### Osmometer measurements

The osmolarity of PSY and AG media were measured using a Wescor Vapro 5520 vapor pressure osmometer. Before use, the osmometer was calibrated using 100 mM, 300 mM, and 1000 mM OptiMole standards from ELITechGroup.

### *B. diazoefficiens* pregrowth

5 mL of fresh media was inoculated with multiple colonies per strain were picked using a sterile stick. After 2-3 days, when the cultures were visibly turbid, these cultures were subcultured into 5 mL of fresh media and allowed to grow to mid-late exponential phase (OD_600_ 0.5-1). The subculturing was repeated, and the second subculture was used as the inoculum for all experiments unless otherwise noted.

### Growth curves

The growth curve assays were performed in 96 well tissue culture plates (Genesee Scientific) using a Spark 10M multimode microplate reader (Tecan, Grödig, Austria). Wells were topped with 50 μL autoclaved mineral oil. Optical density absorbance was taken at 600 nm at 30-minute increments at 30°C with continuous linear shaking.

### Growth curve parameter estimation

To estimate the maximum specific growth rate (μ_m_) and lag time (λ) for the growth curves, the data was fit using an R application that relies on nonlinear least squares to fit nonlinear models to the following Gompertz curve equation (45, 46):

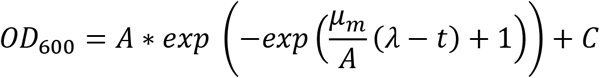

where A is the final OD_600_, t is the time in hours, and C is an adjustment for initial OD_600_. The R application can be found at the following github repository: https://github.com/scott-saunders/growth_curve_fitting The specific version of the growth fitting R application used was retrieved on July 13, 2021 and can be found here: https://github.com/scott-saunders/growth_curve_fitting/blob/master/growth_curve_fitting_ver0.2.Rmd This is a direct link to the application that can be run locally: https://scott-h-saunders.shinyapps.io/gompertz_fitting_0v2/#section-parameter-estimates

### Viable-cell plate counts

Viable-cell plate counts were performed by serially diluting samples in fresh AG media. Dilutions spanning 6 orders of magnitude were plated on AG agar plates as 10 μl drips. Plates were incubated at 30°C. Colonies were counted after 4 days for WT and Δ*hpnH* complement and after 5 days for Δ*hpnH*.

### Lipid packing

Three biological replicates were grown at 30°C and harvested at mid-exponential (OD_600_ = 0.5). Cells were kept at 30°C throughout the entire procedure. Harvested cells were washed 2x by pelleting at 5000 rcf for 7 minutes and resuspending in fresh medium at an OD_600_ of 0.2. Cells were then transferred to black bottom 96-well plates and stained with 80 nM Di-4 ANEPPDHQ (ThermoFisher, D36802). All spectroscopic measurements were carried out using a SPARK 20 plate reader (Tecan, Grödig, Austria) equipped with a thermostat capable of maintaining the temperature with the accuracy of ± 1°C. Samples were incubated for 30 minutes at 150 rpm. Fluorescence emission was measured in the ‘top reading mode’ of the setup in the epi-configuration using a 50/50 mirror and two monochromators (for selecting excitation and emission wavelengths). The sample was excited with xenon flash lamp with the excitation monochromator set to 485 nm (20 nm bandwidth). The fluorescence emission was measured at 540 nm and 670 nm wavelength (20 nm bandwidth each).

General polarization (GP) was calculated from Di4 emission using following formula:

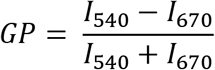

where *I* is the fluorescence emission intensity at given wavelength after a subtraction of the signal measured for the blank suspension.

### Fluorescein diacetate (FDA) permeability assay

To assess relative changes in permeability we employed an assay based on the hydrolysis of FDA to fluorescein after it diffuses into the cell (47, 48). Three biological replicates of each strain were grown until till early exponential (OD_600_ = 0.5). Harvested cells were washed 2x by pelleting at 5000 rcf for 7 minutes and resuspending in fresh medium at an OD_600_ of 0.2. Cells were then transferred to black bottom 96-well plates and FDA (Sigma F7378) was added directly to each well to a final concentration of 20μM. All spectroscopic measurements were carried out using a SPARK 20 plate reader (Tecan, Grödig, Austria) equipped with a thermostat capable of maintaining the temperature with the accuracy of ± 1°C. Immediately following the addition of FDA, samples were measured every 3 minutes for 39 minutes. Excitation was set to: 485 nm (20 nm bandwidth), and emission intensity was measured at 525nm (20 nm bandwidth). The relative permeability of FDA hydrolysis was estimated as the slope of fluorescence vs. time for 39 minutes.

## Results

### Hopanoids are conditionally essential in *B. diazoefficiens*

Due to the previous difficulties isolating a *shc* deletion strain in *B. diazoefficiens*, we decided to construct a conditional SHC expression strain in a Δ*shc* background. The conditional expression system employed a cumate repressor system that was modified for *B. diazoefficiens*, where cumate relieves repression of transcription of the gene of interest (Table S1). With the cumate conditional SHC expression strain in hand, we tested the growth of the strain on plates made up of different media commonly used to cultivate rhizobia *in vitro* (Figure 2). The cumate conditional SHC expression strain was unable to grow on PSY media without cumate present. Therefore, in PSY media, hopanoid production appears to be essential for the growth of *B. diazoefficiens*. However, when we tested for growth on AG medium plates, we observed moderate growth compared to WT without cumate added. This unexpected result indicated that hopanoids are not essential under this condition. With this in mind, we considered the compositional differences between these two media (Table 1). While there are many altered components between the two media, two aspects that stood out to us were the differences in osmolarity and the differences in divalent cation concentrations. Osmolarity is thought to be elevated in the root nodule and can span a wide range in the soil environment (9). Divalent cations are known to stabilize the outer membrane through interactions with LPS, especially calcium (23, 49). The medium osmolarity was 16 mM greater and the divalent cation concentration was almost doubled in PSY compared to the AG medium (Table 1), suggesting that hopanoids are necessary to withstand certain levels of osmolytes and/or ionic strength in *B. diazoefficens*.

**Table 1.** Differences between PSY and AG Media

| Medium | Buffer | Divalent Cations | Carbon Sources | Osmolarity | pH |
| --- | --- | --- | --- | --- | --- |
| PSY | 4.3 mM Phosphate | 45 mM $\text{CaCl}_2$<br>400 mM $\text{MgSO}_4$ | Arabinose<br>Yeast Extract<br>Peptone | $53 \pm 0.06$ mOsM | 7 |
| AG | 5.5 mM HEPES<br>5.6 mM MES | 90 mM $\text{CaCl}_2$<br>730 mM $\text{MgSO}_4$ | Arabinose<br>Yeast Extract<br>Sodium Gluconate | $69 \pm 1.2$ mOsM | 6.6 |

**Figure 2.**
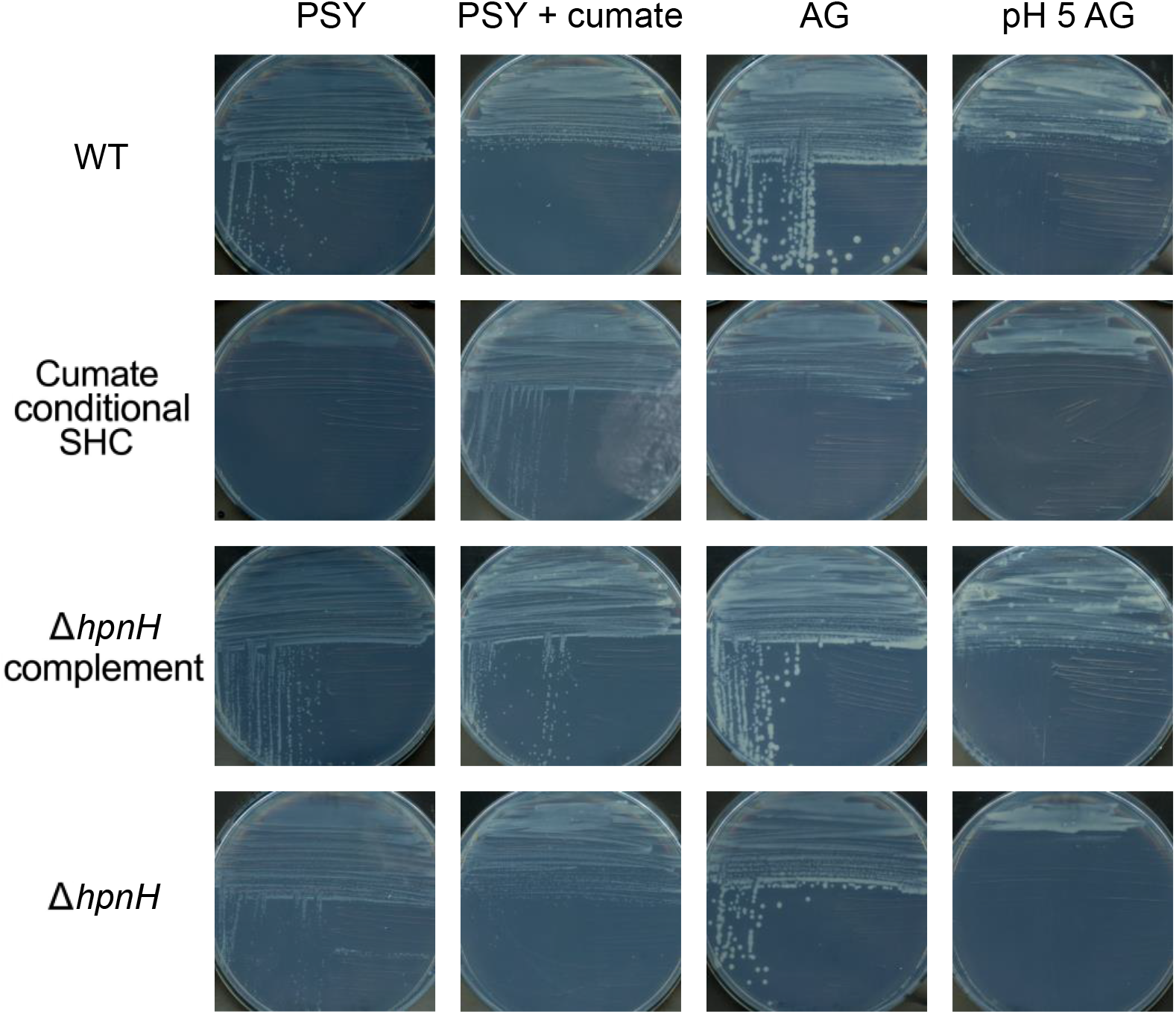
Hopanoids are conditionally essential in *B. diazoefficiens.* A Δ*shc* deletion mutant with a chromosomally integrated cumate-inducible *shc* gene (cumate conditional SHC) is unable to grow on a PSY medium agar plate, but this strain can grow on a PSY agar plate containing cumate which restores hopanoid production. On an AG medium agar plate, the cumate conditional SHC mutant can grow more than on the PSY, but less growth than on PSY with cumate. There is visually greater growth on pH 5 AG medium than on PSY. WT and the Δ*hpnH* complement strains can grow in all conditions. The Δ*hpnH* strain can grow in all conditions as well, with the best growth on the AG medium agar plate.

Despite this realization, we continued to struggle with obtaining a clean *shc* deletion mutant in AG medium, likely due to the fact that the sucrose-selection method we were using to generate the mutant strain (50) exposed it to an osmotic stress beyond the threshold it could tolerate. Accordingly, to probe how the medium composition affects *B. diazoefficiens* strains lacking hopanoids and what this reveals about the ecophysiological role of hopanoids more generally, we turned to the *B. diazoefficiens* Δ*hpnH* strain. To verify that the experimental results using the *B. diazoefficiens* Δ*hpnH* strain are due to the lack of extended hopanoids, we constructed a complement strain with the *hpnH* gene under control of the P*_aphII_* constitutive promoter integrated at the *scoI* locus, as used previously to express a range of fluorescent proteins (44). These strains were tested under the same conditions, and we observed that Δ*hpnH* also showed better growth on AG medium plates compared to PSY plates, while the Δ*hpnH* complement strain behaved similarly to WT. Previously, Δ*hpnH* was shown to be unable to grow at pH 6 in PSY medium (25). The pH within the root nodules and in the legume rhizosphere is known to be acidic (7, 51), so we tested the growth of our strains on pH 5 AG media plates (Figure 2). Interestingly, while growth was lower for the Δ*hpnH* strain and the *shc* conditional mutant at this pH, they were still able to grow. This result indicates that hopanoids are not essential at low pH under all conditions.

### Extended hopanoids protect *B. diazoefficiens* in stationary phase and at low pH

To better understand how the Δ*hpnH* strain grows in AG medium, we quantified different aspects of the growth curve. In PSY, the Δ*hpnH* strain has a pronounced defect in both exponential and stationary phase (25). Even when Δ*hpnH* and WT cultures were sampled at the same OD_600_ in exponential phase, Δ*hpnH* had drastically lower viability than WT; we reasoned this might be due to initial inoculum viability differences from “overnight” cultures. To test and pre-empt this, we subcultured twice from an initial turbid culture inoculated with colonies from a fresh plate. Using this technique, we observed that the Δ*hpnH* mutant strain has only a very slight growth defect compared to WT and the Δ*hpnH* complement in the pH 6.6 AG medium (Figure 3). To confirm that the similarity in OD_600_ is due to a comparable number of viable cells, we determined colony forming units (CFUs) after 24 hours (mid-late exponential) and 72 hours (stationary phase) of growth. After 24 hours, the CFUs were very similar but after 72 hours, the Δ*hpnH* strain was much worse off, leading to a significant difference in CFUs between the WT and Δ*hpnH* strain. This result confirms the stationary phase defect observed previously in PSY medium, and that our culturing method successfully removes differences in inocula that influenced previous experiments. Using this approach, we tested the Δ*hpnH* strain’s growth in the pH 5 AG medium (Figure 3 and Figure S1). Here, we observed a significant defect in growth rate and increased stationary phase death for the Δ*hpnH* strain compared to WT and the Δ*hpnH* complement. Upon inspection, clumping of the Δ*hpnH* strain was observed in the wells of the plate, perhaps indicating death followed by increased biofilm formation.

**Figure 3.**
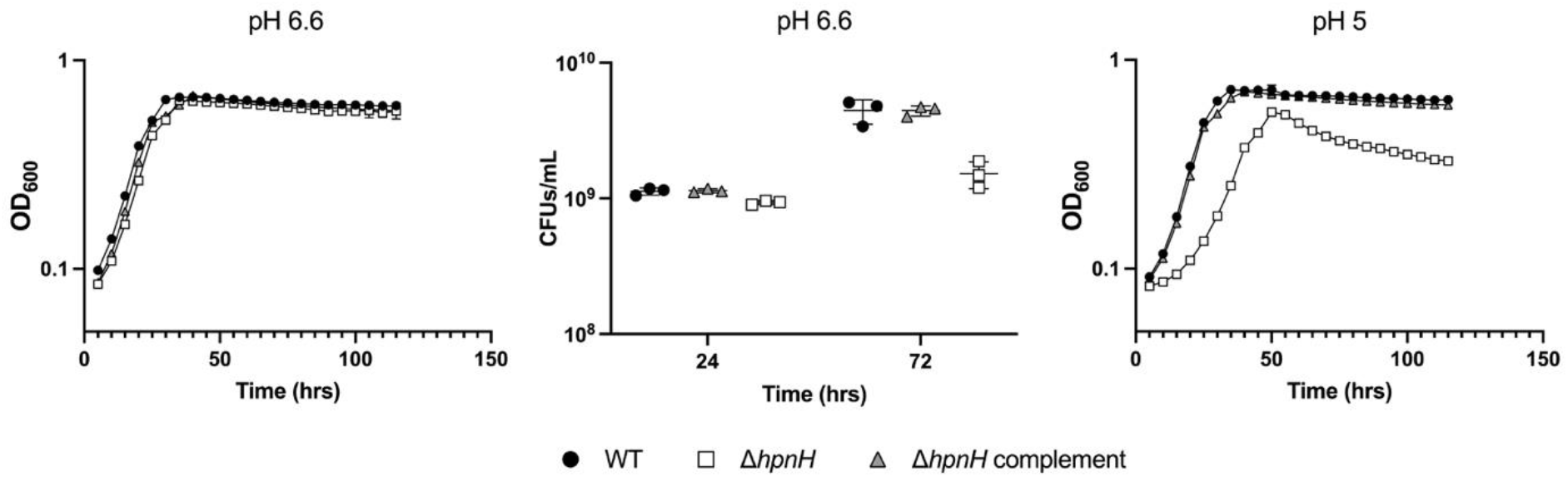
*B. diazoefficiens* Δ*hpnH* strain is sensitive to low pH and stationary phase. (A, C) Growth of WT (black circles), Δ*hpnH* complement (grey triangles), and Δ*hpnH* (white squares) in AG media at pH 6.6 and pH 5 was monitored at an optical density of 600 nm (OD_600_). Each curve represents the average of three biological replicates. (B) Colony forming units per mL (CFUs/mL) were measured for WT (black circles), Δ*hpnH* complement (grey triangles), and Δ*hpnH* (white squares) strains grown in AG media at pH 6.6 during exponential phase (24 hrs) and stationary phase (72 hrs). Error bars (standard deviation) are included (A-C), but some are smaller than the point markers.

### Physicochemical medium conditions affect the growth of *B. diazoefficiens* Δ*hpnH*

Having discovered that growth defects are condition-dependent for the hopanoid deficient μ*shc* strain, we decided to check whether this was also true for the growth defect of the Δ*hpnH* strain in pH 5 AG medium. First, we tested the effects of osmolarity by adding the nonmetabolizable, nonionic osmolyte inositol (Figure 4). The growth rate and lag time parameters were estimated using a Gompertz model. As the concentration of inositol was increased from 25 mM to 400 mM, the growth rate decreased for WT and the Δ*hpnH* complement (Figure S2). However, the growth rate for the Δ*hpnH* strain increased to a maximum growth rate with 100 mM inositol added, before decreasing. The Δ*hpnH* strain never reached the same growth rate as WT, but the differences in the growth rate response over this osmolarity range illustrates that the strains experience these conditions very differently and that a “Goldilocks” osmotic zone exists for the Δ*hpnH* mutant where its growth is enhanced. A similar trend was observed with lag time, where the Δ*hpnH* strain exhibited a minimum lag time at 100 mM inositol added. The lag time of the WT and the Δ*hpnH* complement remained relatively constant up to 100 mM inositol added and then steadily increased. These experiments were also completed with sorbitol as the osmolyte for WT and the Δ*hpnH* strain, showing similar results (Figure S3).

**Figure 4.**
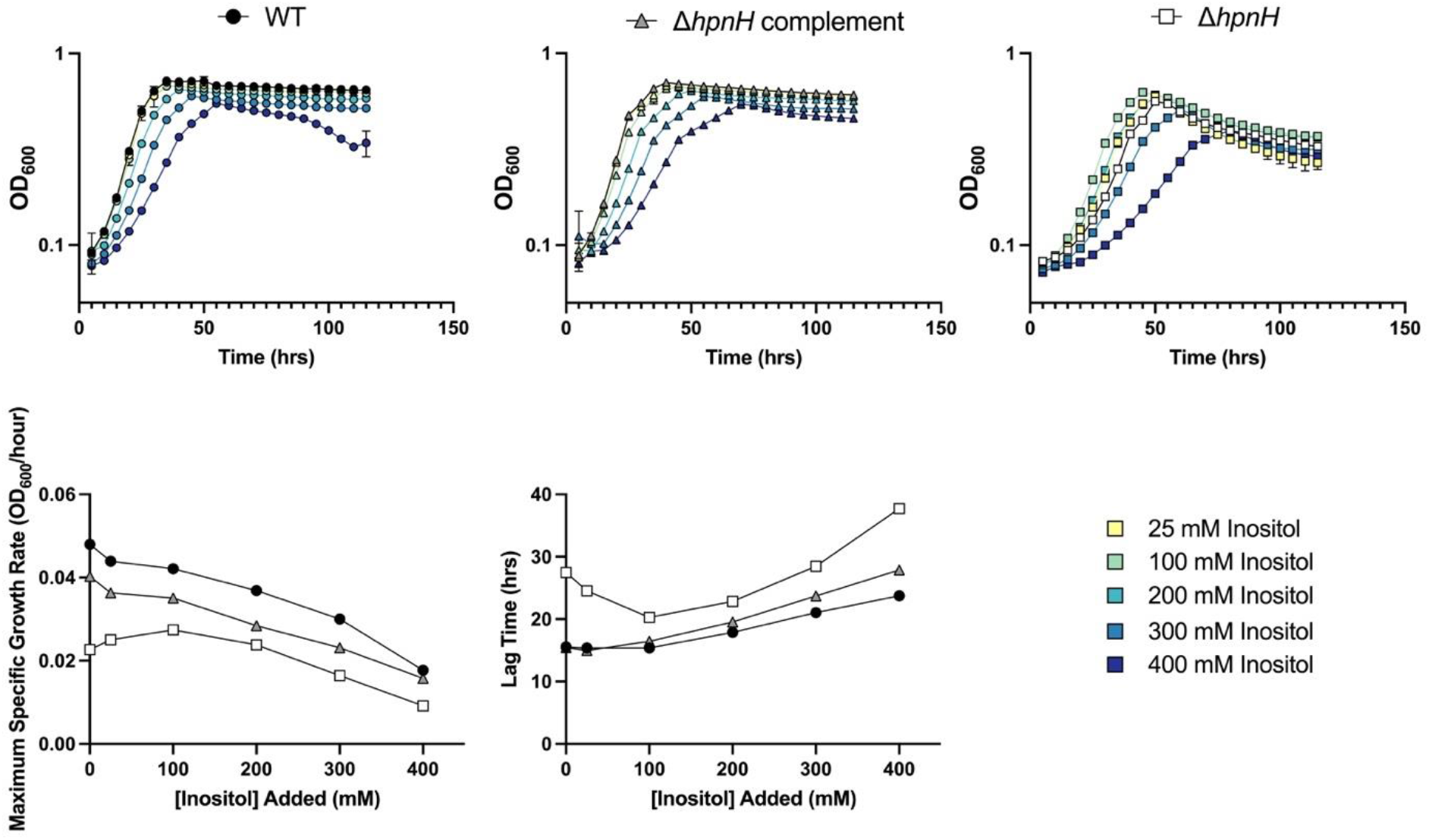
*B. diazoefficiens* Δ*hpnH* strain growth is sensitive to the concentration of inositol. (A) Growth of WT (black circles), Δ*hpnH* complement (grey triangles), and Δ*hpnH* (white squares) in AG media at pH 5 with increasing concentration of inositol was monitored at OD_600_. The colors of the markers correspond to different concentrations of inositol as noted in the legend. (B) Growth rate (μ) and lag time were quantified by fitting a single Gompertz to each growth curves from (A). The results are plotted according to increasing concentration of inositol. Arrows point to the concentration of inositol (100 mM) where the growth of the Δ*hpnH* strain is optimized. Error bars (standard deviation) are included, but some are smaller than the point markers.

Next, we tested the effects of divalent cations on growth. Because the AG medium already has almost double the divalent cation concentration as the PSY medium, we created a “low dication” pH 5 AG medium using the PSY concentrations of divalent cations, specifically magnesium and calcium (45 mM Ca^2+^ and 400 mM Mg^2+^). We then increased the calcium ion concentration in this low dication AG medium. The growth rate and lag time for WT and the Δ*hpnH* complement were agnostic to these changing conditions (Figure 5). However, the Δ*hpnH* strain grew at a slower rate and with a longer lag time in the low dication AG medium condition, indicating that 45 μM may be particularly stressful for the Δ*hpnH* strain. The growth rate recovered with 100 μM additional calcium and remained at the same growth rate. The lag time decreased as well as the calcium concentration was increased. When 500 μM additional calcium was added to the low divalent cation condition, the growth curve for Δ*hpnH* became almost indistinguishable from the pH 5 AG medium. These experiments were also completed with magnesium as the divalent cation for WT and the Δ*hpnH* strain with similar but less pronounced results (Figure S4).

**Figure 5.**
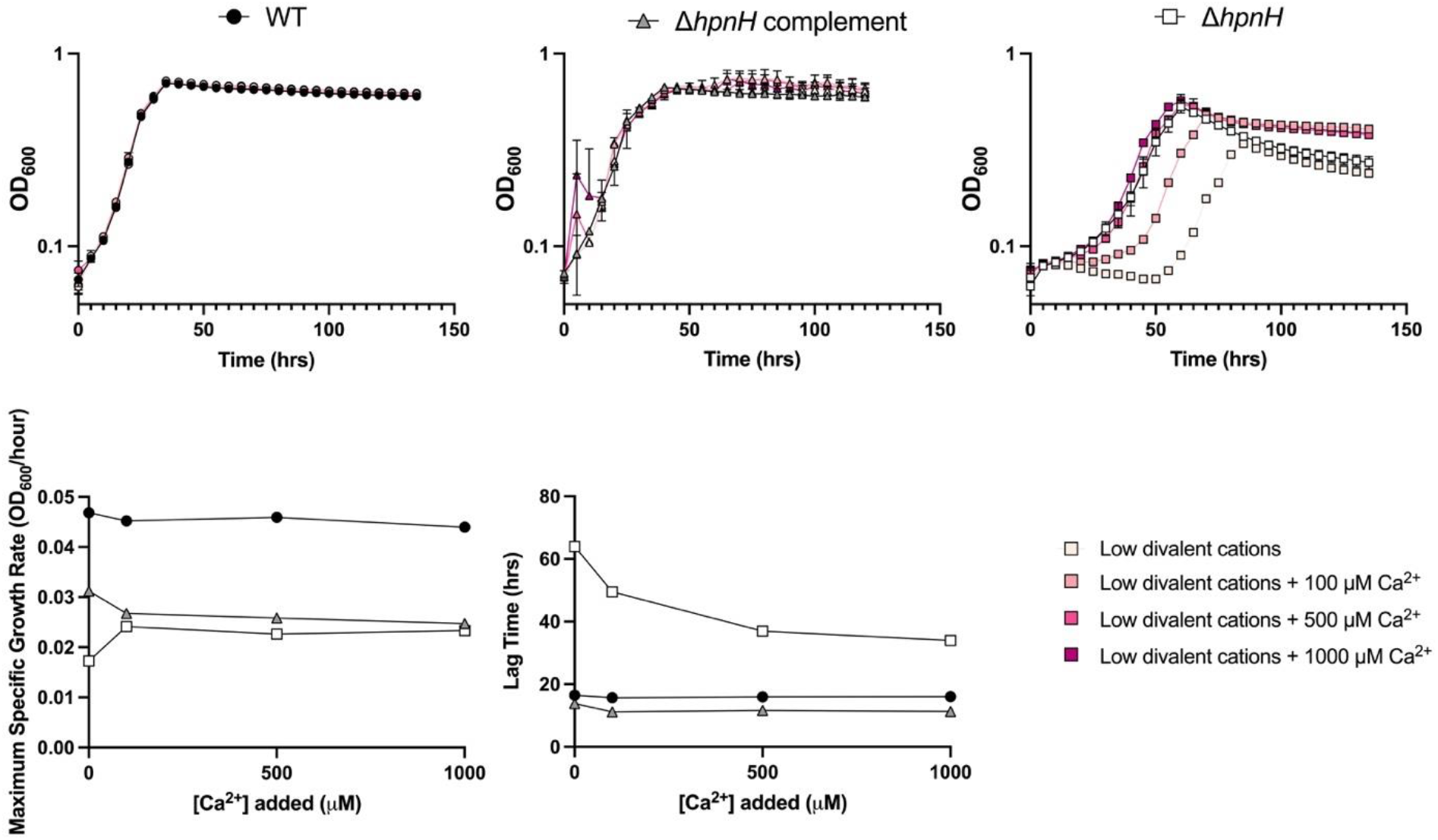
*B. diazoefficiens* Δ*hpnH* strain growth is sensitive to the concentration of calcium. (A) Growth of WT (black circles), Δ*hpnH* complement (grey triangles), and Δ*hpnH* (white squares) in AG media at pH 5 with different concentrations of divalent cations was monitored at OD_600_. The colors of the markers correspond to different concentrations of Ca^2+^ ions as noted in the legend. (D) Growth rate (μ) and lag time were quantified by fitting a single Gompertz curve to each growth curve from (C). The results are plotted according to increasing concentration of Ca^2+^ ions with the low dication condition included at y=0. All growth curves and quantifications represent the average of three biological replicates. Error bars (standard deviation) are included, but some are smaller than the point markers.

Together, these results indicate that the lack of extended hopanoids can be compensated for by changing the physicochemical properties of the growth medium, specifically the osmolarity and divalent cation concentration. Unlike the pattern seen for osmolytes for Δ*hpnH*, where an intermediate concentration minimized lag time and increased growth rate, increasing concentrations of divalent cations increasingly shrunk the lag time yet did not appreciably affect growth rate.

### Extended hopanoids are required to regulate the membrane properties of *B. diazoefficiens*

In order to explain the growth phenotypes of the Δ*hpnH* strain, we hypothesized that extended hopanoids play a role in buffering the outer membrane against physicochemical perturbations. Indeed, the unextended hopanoid, diplopterol, is known to modulate changes in lipid A packing *in vitro* that occur in response to decreased pH (52). To probe the underlying mechanism behind the physicochemical compensation for loss of extended hopanoids, we used the lipophilic dye Di-4-ANEPPDHQ (Di-4) (Figure 6A). The general polarization (GP) of Di-4 reports on lipid packing, with higher GPs indicative of increased packing. Because lipid packing is correlated with viscosity and bilayer stability it provides a robust general indicator of changes in membrane biophysical properties (53, 54). Because of its size (665.55 MW) and polarity, Di-4 should preferentially label the outer leaflet of the outer membrane, and has been routinely used to monitor changes in surface membrane lipid packing (55). However, given the interplay between membrane stability and permeability, the assumption that Di-4 selectively labels the outer leaflet of the outer membrane may not hold if membrane permeability is increased sufficiently to allow Di-4 to cross the outer membrane. Outer membranes contain saturated Lipid A and have been shown to have similar lipid packing to liquid ordered phase membranes in vitro (56) whereas inner membranes comprise more disordered phospholipids, which should have lower lipid packing similar to a liquid disordered phase. Thus, a large decrease in Di4 GP could either be interpreted as a decrease in outer membrane lipid packing, or to a large increase in outer membrane permeability allowing Di4 to label the inner membrane. Both results would indicate a large change in the mechanical properties of the outer membrane.

**Figure 6.**
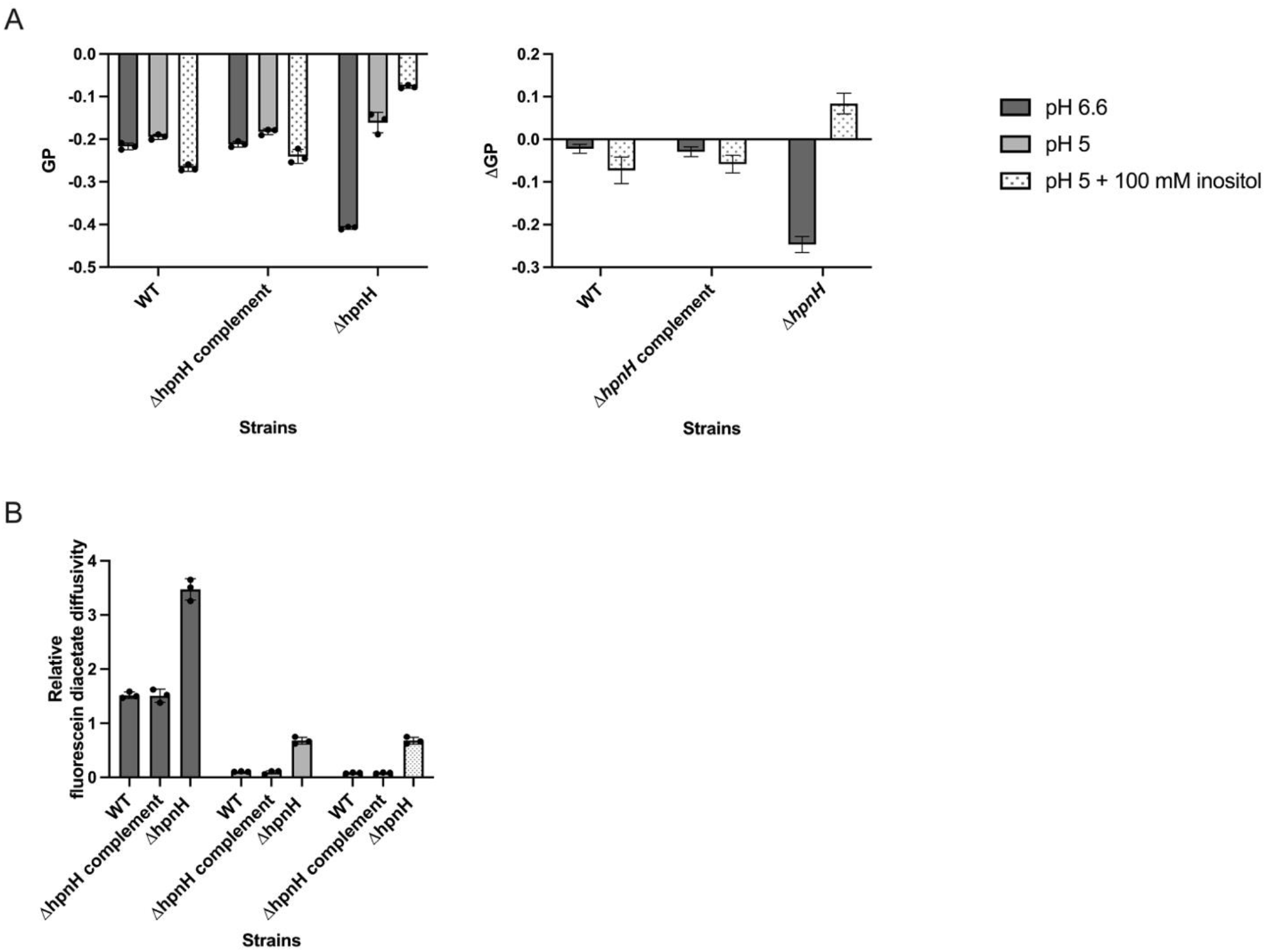
*B. diazoefficiens* Δ*hpnH* strain is deficient in its ability to regulate its membrane properties. (A) Lipid packing measured by Di-4 general polarization (GP) index for WT, Δ*hpnH* complement, and Δ*hpnH* strains grown in AG media with pH 6.6 (dark grey), pH 5 (light grey), and pH 5 with 100 mM added inositol (grey dots). Individual measurements are shown as black circles. ΔGP for each strain compared to the AG pH 5 condition. Error bars (standard deviation) are included. (B) Relative cell envelope permeability measured by a fluorescein diacetate diffusivity assay of WT, Δ*hpnH* complement, and Δ*hpnH* strains grown in AG media with pH 6.6 (dark grey), pH 5 (light grey), and pH 5 with 100 mM added inositol (grey dots). Individual measurements are shown as black circles. Error bars (standard deviation) are included.

To evaluate the role of extended hopanoids in outer membrane acclimation to pH and osmotic strength, we compared Di-4 GP in cells grown at pH 6.6 and 5, and cells grown at pH 5 in the presence and absence of inositol. The Di-4 GP values for WT and the Δ*hpnH* complement strains were almost identical across the three conditions (pH 6.6 AG media, pH 5 AG media, and pH 5 AG media + 100 mM inositol), which indicates that the outer membrane of our complement strain is responding very similarly to WT. The Δ*hpnH* strain had lower GP values than the WT and the Δ*hpnH* complement when grown in pH 6.6 AG media. Additionally, all three strains have higher GP values when grown in pH 5 AG medium, which is consistent with observations that low pH increases the packing and order of LPS (52, 57). Interestingly, WT and the Δ*hpnH* complement showed a small decrease in GP values in AG pH 5 medium with 100 mM inositol added, while the Δ*hpnH* strain GP values continued to increase.

To better interpret these results, we determined the ΔGP values, comparing the change in GP compared to the standard pH 6.6 AG medium condition. WT and the Δ*hpnH* complement underwent very small changes compared to the Δ*hpnH* strain. In addition to exhibiting comparably more GP variability with changing osmolarity, Δ*hpnH* showed a very large negative shift of around 0.3 GP at pH 6.6. Such a large negative shift in GP suggested a considerable change in mechanical properties of the outer membrane at higher pH. To determine whether this might be the result of compromised outer membrane integrity, we examined whether cell envelope permeability of Δ*hpnH* at pH 6.6 was higher than for other conditions and strains. We estimated changes in membrane permeability using an assay based on relative changes in fluorescein diacetate (FDA) diffusivity (47). FDA is non-fluorescent, and rapidly diffuses into the cell where it is hydrolysed. Fluorescein, which is fluorescent, is produced from the hydrolysis of FDA. Additionally, because fluorescein is charged, it cannot diffuse rapidly out of the cell. Therefore, the rate of increase in fluorescein fluorescence can provide an estimate of the relative permeability of FDA across the cell envelope of different strains or across varying growth conditions. We observed that Δ*hpnH* had nearly 3-fold higher permeability than WT at pH 6.6, and nearly 10-fold higher permeability compared with Δ*hpnH* at pH 5 with and without inositol (Figure 6B). The large negative shift in Di-4 GP values at pH 6.6, therefore is consistent with both reduced lipid packing and compromised membrane integrity. Overall, these results show that extended hopanoids play an important role in buffering *B. diazoefficiens* membrane properties against physicochemical perturbations.

## Discussion

By making a serendipitous observation of the conditional essentiality of the *shc* gene in *B. diazoefficiens*, we discovered that the physicochemical environment is extremely important for the growth of hopanoid deficient *B. diazoefficiens* strains, including the extended hopanoid deficient mutant, Δ*hpnH*. We confirmed that the Δ*hpnH* strain undergoes significant death in stationary phase, but that, in contrast to previous results (25), it can grow at pH 5 in a medium with higher osmolarity and divalent ion concentration. We identified environmentally relevant conditions that partially compensate for the *B. diazoefficiens* Δ*hpnH* mutant growth defect at low pH: intermediate osmolarity and elevated divalent cation concentrations. Finally, using a biophysical technique, we discovered that extended hopanoids are important for modulating lipid packing of the outer membrane.

When we first discovered that our conditional Δ*shc* mutant could grow on solid AG medium without cumate induced hopanoid production but not on PSY medium, it caused us to reexamine our previous results with Δ*hpnH* carried out in PSY (25). As previously shown, Δ*hpnH* has a stationary phase defect and increased lag time when grown in PSY. When additional stress was added to the PSY condition, such as increased temperature, lowered pH, or microoxia, the Δ*hpnH* strain was unable to grow at all. It is possible that due to its stationary phase defect under these conditions, the Δ*hpnH* inoculum in these experiments may have had fewer viable cells than WT, despite similar optical density measurements, contributing to the severity of these phenotypes. Indeed, the “control” growth of the Δ*hpnH* mutant is visually reduced compared to the WT in the stressor gradient plate assay performed in these studies, consistent with this hypothesis. With this in mind, our revised inoculation protocol enabled us to see that Δ*hpnH* grows very similarly to WT in AG medium and can even grow at pH 5. Surprisingly, Δ*hpnH* experiences significant death in stationary phase compared to WT in AG medium, despite the optical density measurements remaining constant. These results highlight the importance of not relying on optical density measurements to assess bacterial viability, as they can be de-coupled; a known but often forgotten phenomenon. It is likely that the previous results in PSY were accentuated due to differences in inoculum viability, representing the combined effects of the stationary and exponential phase growth defects.

As we tried to understand why the two media affect the growth of our hopanoid deficient mutants differently, we reflected on their composition. Three differences stood out to us: differences in pH, divalent cations, and osmolarity. Previous research on Δ*shc* mutants in other closely related bacteria have shown that hopanoids are important in both acidic and basic conditions. Specifically, a Δ*shc* mutant of *Rhodopseudomonas palustris* failed to grow as it made the medium more basic (58). We know that in both PSY and AG media, *B. diazoefficiens* increases the pH through amino acid metabolism of the complex media (i.e., yeast extract). However, the AG medium also contains a higher buffering capacity at a lower pH than PSY (6.6 vs 7), thus likely extending the time before the pH is increased substantially. Therefore, the conditional Δ*shc* mutant’s lack of growth on PSY medium and growth on AG medium is perhaps not entirely surprising and illustrates how important medium composition may be when isolating and growing a mutant in hopanoid production or other membrane component.

The higher concentration of divalent cations, specifically, magnesium and calcium, in the AG medium compared to PSY, was interesting because of the role these cations play in outer membrane cohesion. Specifically, magnesium and calcium intercalate within the LPS layer, shielding negatively charged residues, resulting in a more ordered and robust outer membrane in the face of physicochemical stressors (23, 49). Our two hopanoid deficient mutants cannot make HoLA, a component of LPS that contributes to membrane ordering (27, 59). Recent work in *Bradyrhizobium* BTai1 has revealed that calcium ions increase membrane bilayer thickness, an indication of increased membrane order, in membrane vesicles containing LPS without a hopanoid attached (59); this phenomenon suggests a mechanism whereby calcium may be able to compensate for lack of hopanoids. Our work shows that the lag time of Δ*hpnH* in pH 5 AG medium can be reduced by increasing concentrations of divalent cations, supporting this hypothesis. Calcium has a greater effect than magnesium, likely reflecting the fact that calcium ions more strongly increase lipid bilayer rigidity through dehydration effects than magnesium ions (60). WT was agnostic to these changes in divalent ion concentration perhaps due to the presence of HoLA, unlike other Bradyrhizobia strains (61). Interestingly, while calcium is maintained at low concentrations (0.1 μM) in the cytosol of plant cells, calcium has been shown to localize to the root nodule (62). Calcium binding proteins were specifically found in the root nodules from *Medicago truncatula* (63). Indeed, sufficient calcium is needed for the bacteria to fix nitrogen in the root nodules (62, 64). This evidence illustrates the importance of calcium within the acidic root nodule (7), and helps rationalize why the Δ*hpnH* <u>mutant grows reasonably well *in planta*:</u> elevated calcium levels may compensate for the loss of extended hopanoids within the root nodule.

We were surprised when we found that the AG medium has a higher osmolarity than the PSY medium, since our previous *in vitro* studies in PSY medium indicated that hopanoids can be protective against hyperosmotic stress (25). Yet, as previously noted, these media are compositionally different in more than one way. It thus appeared possible that, at a lower pH, the relationship between osmolarity and hopanoids might be more nuanced. We hypothesized that hypoosmolarity might also be stressful for hopanoid-deficient mutants that have less robust membranes, and our findings bore this out. A potential mechanism that explains this observation follows: When first introduced to hypoosmotic conditions, water tries to move into the higher osmolarity cell, likely causing at least a transient increase in membrane fluidity as the cell stretches to accommodate the increased volume (65). The cell responds by opening mechanosensitive channels to eject solutes and lower the cytosolic osmolarity while synthesizing osmoregulated periplasmic glucans, thus osmotically buffering the cytoplasm (66). In a cell with a less robust membrane and perhaps increased permeability due to the absence of hopanoids, the increased fluidity during initial water influx may kill some cells, while the periplasmic glucans may be more easily lost to the medium, losing their ability to osmotically buffer the cytoplasm. At low pH, these effects would be magnified as the influx of water would also bring an influx of protons, adding an additional stress. When we added inositol to the pH 5 AG medium, the Δ*hpnH* strain grew better, decreasing the lag time and increasing growth rate up to 100 mM added inositol. Comparatively, WT grew more poorly upon even the smallest addition of inositol (25 mM). This result illustrates that the low osmolarity of the medium is particularly stressful to the Δ*hpnH* strain. Interestingly, inositol makes up a large proportion of the compounds found in the symbiosome space (67). Indeed, the symbiosome space contains approximately 180 mM of low molecular weight compounds (67), notably similar to the maximally restorative osmolarity in our experiments (100 mM inositol pH 5 AG medium). It is thus possible that the root nodule microenvironment allows the Δ*hpnH* strain to survive and fix nitrogen, despite its obvious growth defects at low pH.

Interestingly, while both osmolarity and divalent cation concentrations affect the growth of the Δ*hpnH* strain at low pH, the effects are different. Specifically, divalent cation concentration had the greatest effect on lag time while the added osmolytes affected both lag time and growth rate. These differences suggest that different mechanisms underpin the mutant’s response, despite both having the potential to rigidify the outer membrane. We hypothesize that these differences may arise due to the inositol primarily addressing the root cause of the stress, hypoosmolartiy, while the increase in divalent cations protects against the effects of hypoosmolarity.

The sensitivity of hopanoid-deficient strains to specific external conditions suggested that extended hopanoids might be particularly important in helping cells respond to environmental changes. In contrast to diplopterol, a shorter hopanoid that contains a hydrophilic group, which has been shown to rigidify the membrane while keeping lipids from entering a gel phase and retaining lateral lipid diffusivity (52, 68) extended hopanoids had only been shown to rigidify the membrane (25, 69, 70). Our biophysical experiments confirm that extended hopanoids are necessary for membrane rigidification, but also reveal that lack of extended hopanoids causes greater problems with membrane stability. Δ*hpnH* displays much greater variability in lipid packing between conditions than WT, as evidenced by the larger ΔGP values for the mutant. This result suggests that the WT can adjust its lipid packing to maintain a relatively constant membrane fluidity and is intrinsically more mechanically stable. In contrast, the Δ*hpnH* strain struggles to adjust its lipid packing in response to changes in osmolarity, and membrane integrity is compromised by changes in pH. The lipid packing of the Δ*hpnH* strain is primarily affected by the external environment. In the case of the pH 5 AG medium with 100 mM inositol added, both increased osmolarity and inositol specifically are known to rigidify membranes (65, 71), explaining the increased GP values for the Δ*hpnH* strain. On the other hand, the WT can adjust its membrane to counteract environmentally-triggered membrane rigidification, thus leading to slightly lower GP values. Overall, these results indicate that extended hopanoids play an important role in *B. diazoefficiens* adjustment to and fortification against the external environment.

In conclusion, the lack of hopanoids, and specifically, extended hopanoids—which are required for HoLA biosynthesis—makes *B. diazoefficiens* particularly sensitivity to environmental conditions in ways that are relevant to its lifecycle. That the lack of extended hopanoids can be partially compensated for by a moderately high osmotic level, helps to resolve the paradox of why the Δ*hpnH* mutant can be symbiotically successful if given sufficient time to develop within root nodules. Yet, its sensitivity to hypoosmotic conditions suggest that hopanoids may provide a fitness advantage to rhizobia in waterlogged soils, where osmolytes and divalent cations are diluted. Together, our findings emphasize the importance of considering the full ecophysiological picture when attempting to understand the selective benefits of a given molecular component on an organism. It has been said that the only constant in life is change, a point worth remembering when considering the effects of hopanoids on peripatetic soil organisms.

## Acknowledgements

We thank members of the Newman lab for their helpful comments and insights, especially Brittany Belin and all past members of Team Hopanoid. Thank you to Hans Martin-Fischer for his constant support of our work. This research was enabled by a NSF graduate research fellowship Foundation (E.T.), NASA (NNX16AL96G to D.K.N.), a German Federal Ministry of Education and Research BMBF grant (to J.S., project 03Z22EN12), and a VW Foundation “Life” grant (to J.S., project 93090).

## Figure Legends

**Table S1.**
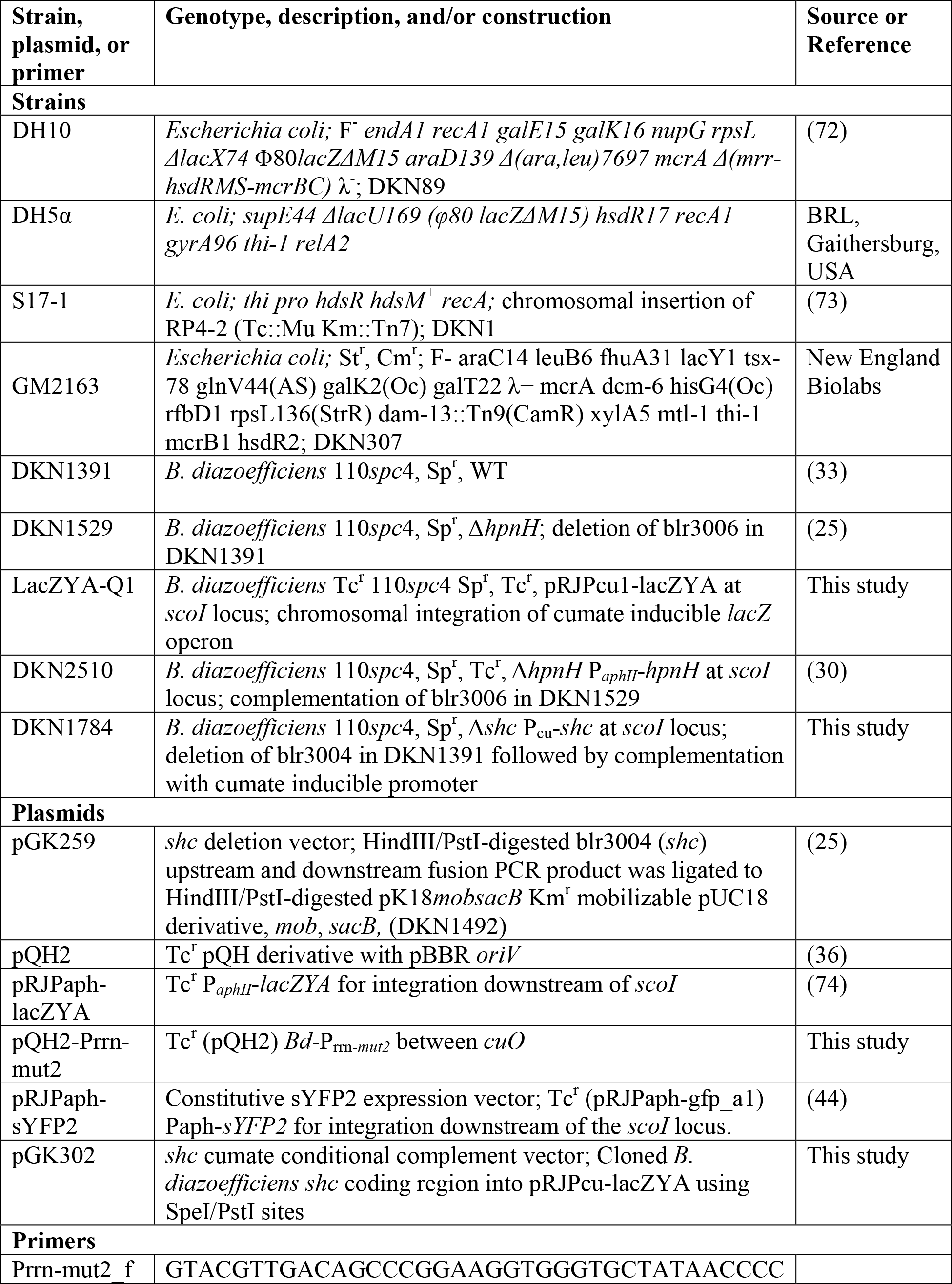
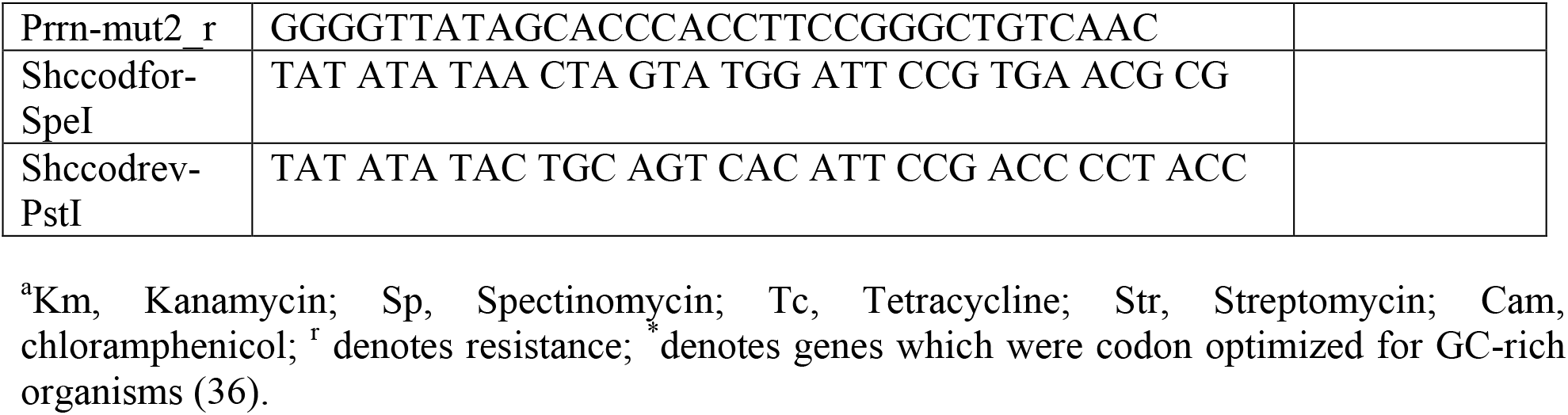
Strains, plasmids, and primers used in this study^a^

**Figure S1.**
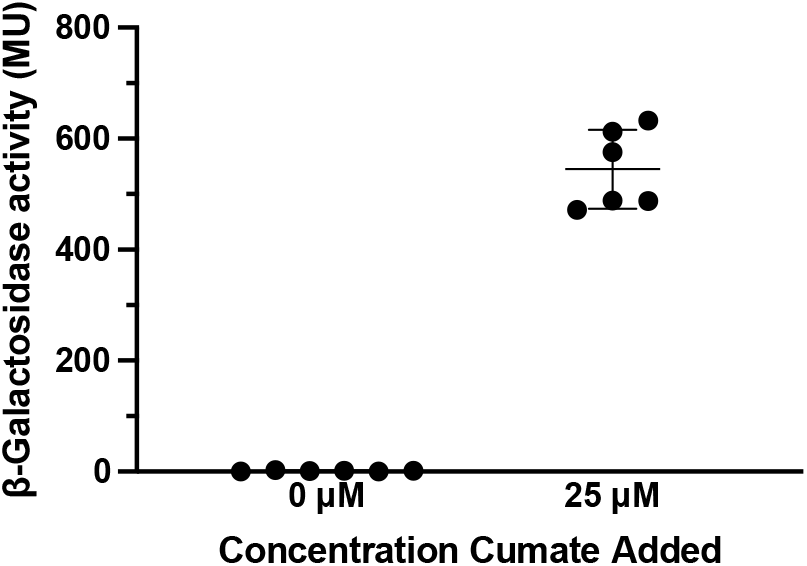
Cumate-inducible system functions as an on/off switch. For the strain LacZYA-Q1, when no cumate inducer is added, there is no β-Galactosidase activity. When 25 μM cumate inducer is added, β-Galactosidase activity is observed.

**Figure S2.**
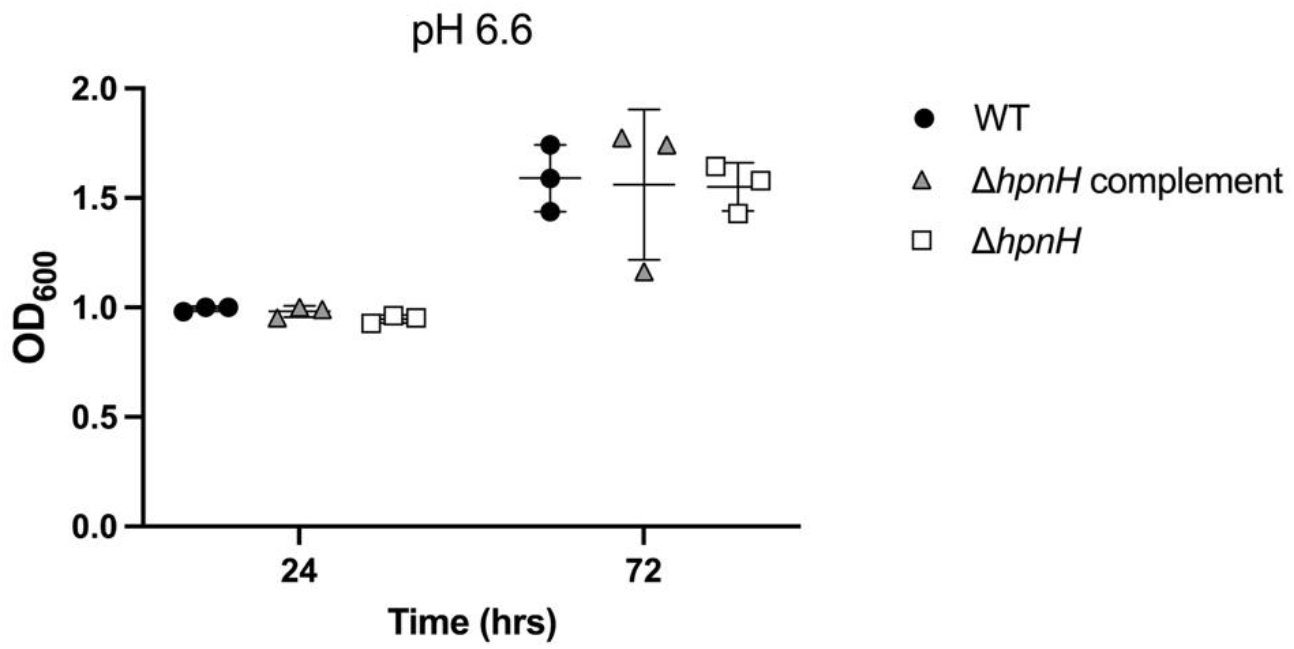
Differences in CFUs/mL not observed by OD_600_. OD_600_ was measured for WT (circles), Δ*hpnH* complement (triangles), and Δ*hpnH* (squares) strains grown in AG media at pH 6.6 during exponential phase (24 hrs) and stationary phase (72 hrs). Error bars (standard deviation) are included, but some are obscured by the point markers.

**Figure S3.**
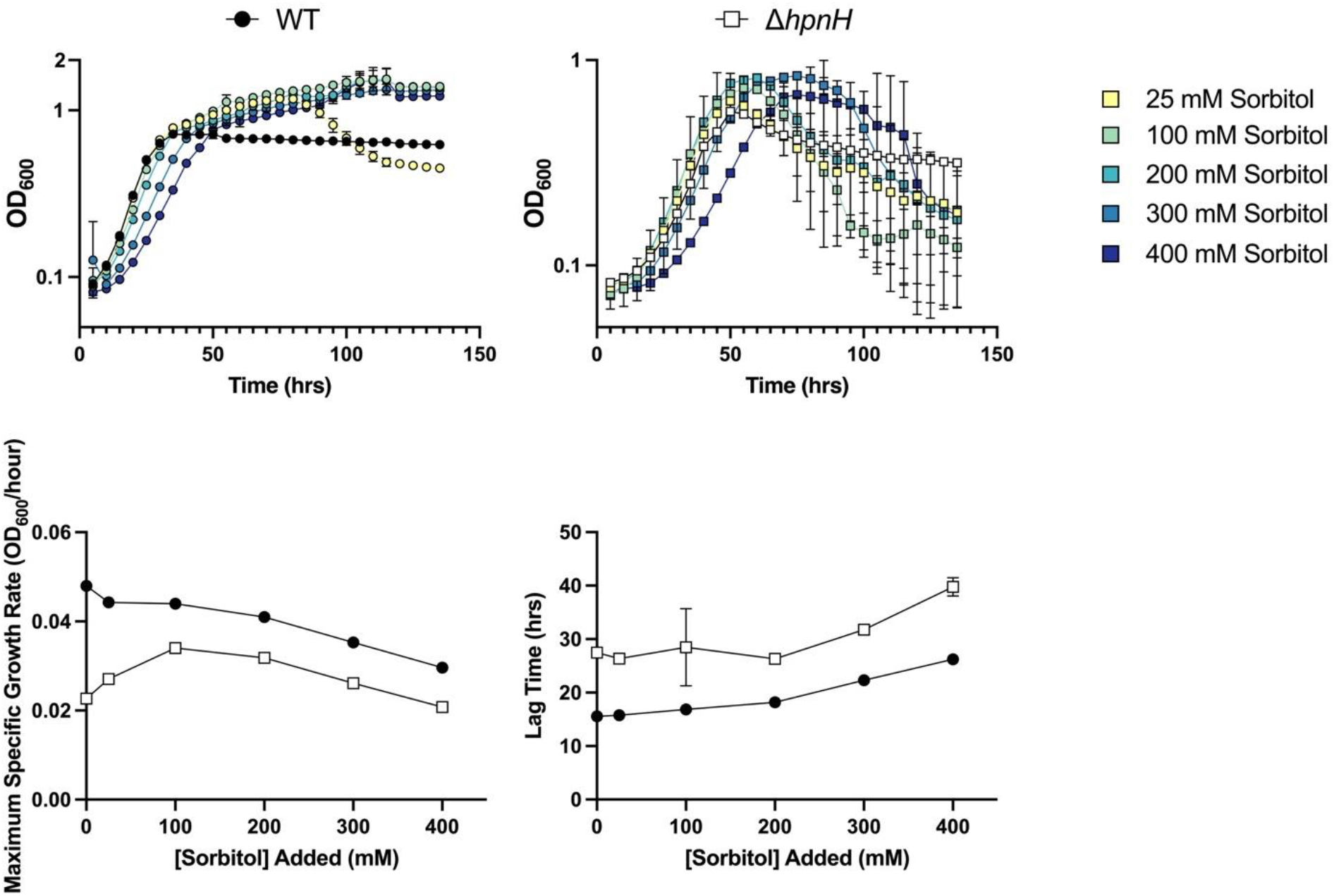
*B. diazoefficiens* Δ*hpnH* strain growth is sensitive to the concentration of sorbitol. (A) Growth of WT (circles), Δ*hpnH* complement (triangles), and Δ*hpnH* (squares) in AG media at pH 5 with increasing concentration of sorbitol was monitored at OD_600_. The colors of the markers correspond to different concentrations of sorbitol as noted in the legend. (B) μ and lag were quantified by fitting a single Gompertz to each growth curve from (A). The results are plotted according to increasing concentration of sorbitol.

**Figure S4.**
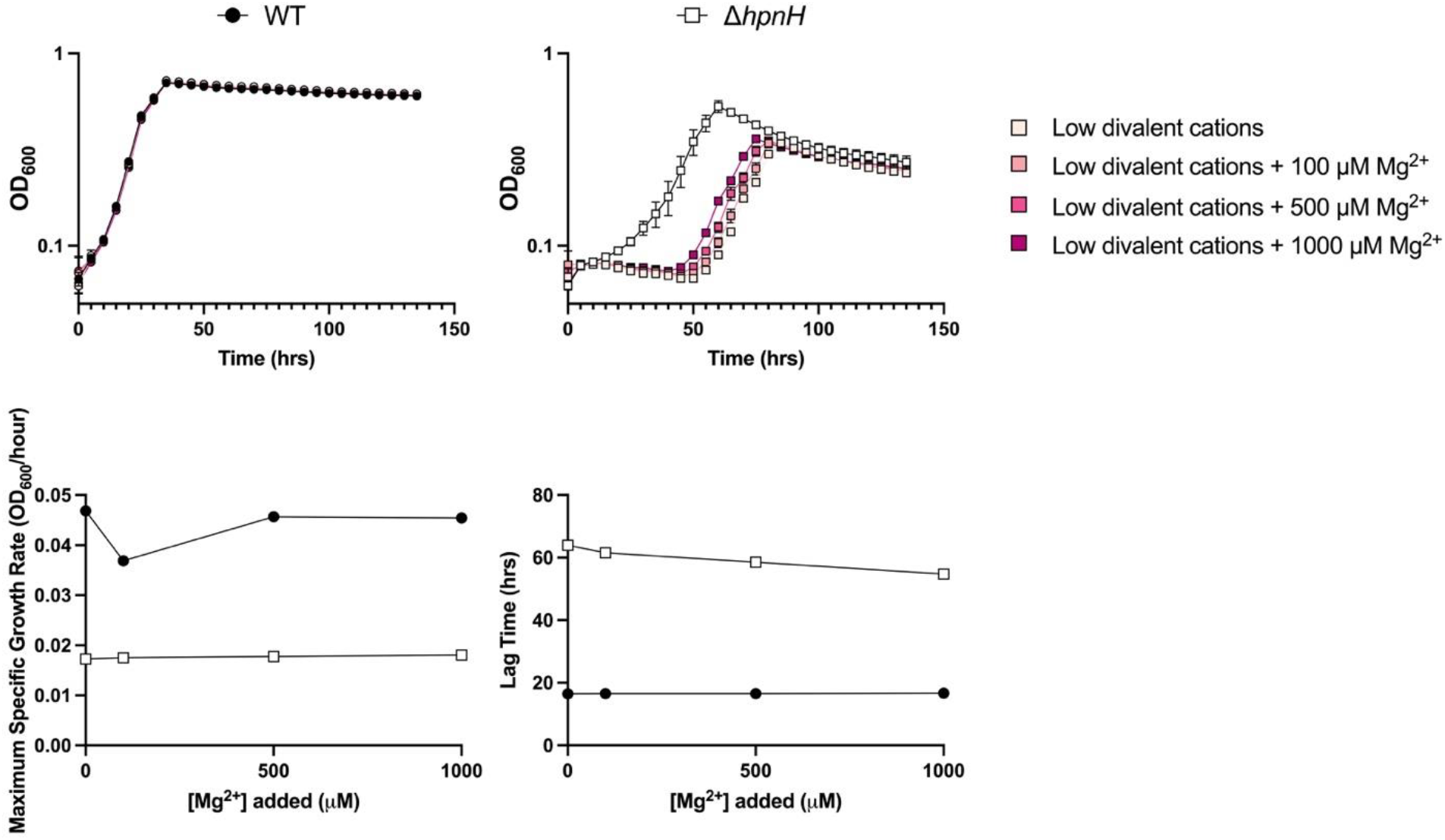
*B. diazoefficiens* Δ*hpnH* strain growth is sensitive to the concentration of magnesium ions. (A) Growth of WT (circles), Δ*hpnH* complement (triangles), and Δ*hpnH* (squares) in AG media at pH 5 with different concentrations of divalent cations was monitored at OD_600_. The colors of the markers correspond to different concentrations of Mg^2+^ ions as noted in the legend. (B) μ and lag were quantified by fitting a single Gompertz curve to each growth curve from (A). The results are plotted according to increasing concentration of Mg^2+^ ions with the low dication condition included at y=0. All growth curves and quantifications represent the average of three biological replicates. Error bars (standard deviation) are included, but some are smaller than the point markers.

**Figure S5.**
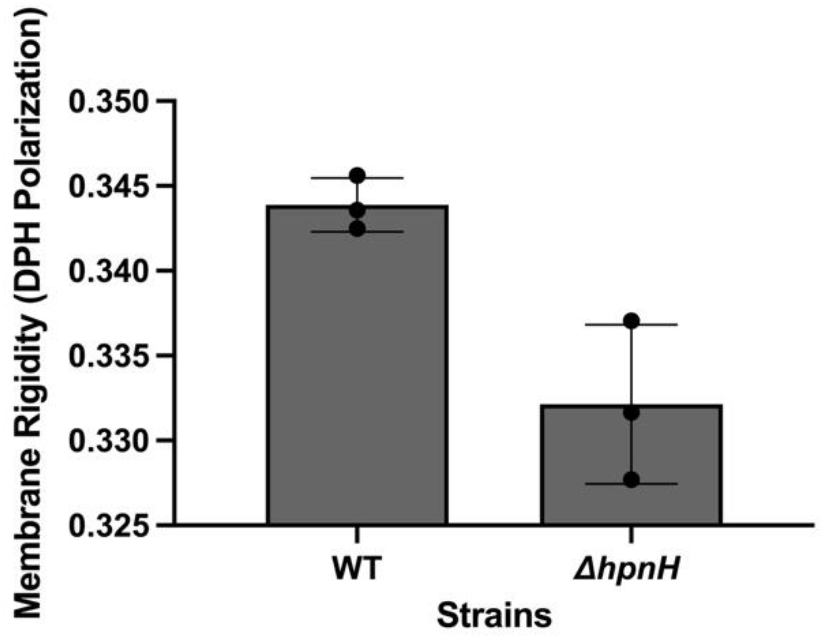
*B. diazoefficiens* Δ*hpnH* strain membrane is less ordered than WT when grown in AG media. Whole-cell membrane fluidity measurements by fluorescence polarization of DPH. Error bars represent the standard deviations from three biological replicates.

## References

1. Doran JW, Zeiss MR. 2000. Soil health and sustainability: managing the biotic component of soil quality. Applied Soil Ecology 15:3–11.

2. Norris CE, Congreves KA. 2018. Alternative management practices improve soil health indices in intensive vegetable cropping systems: A review. Front Environ Sci 6.

3. Tilman D, Cassman KG, Matson PA, Naylor R, Polasky S. 2002. Agricultural sustainability and intensive production practices. Nature 418:671–677.

4. Foyer CH, Nguyen H, Lam H-M. 2019. Legumes-The art and science of environmentally sustainable agriculture. Plant Cell Environ 42:1–5.

5. Oldroyd GED, Murray JD, Poole PS, Downie JA. 2011. The rules of engagement in the legume-rhizobial symbiosis. Annu Rev Genet 45:119–144.

6. Gibson KE, Kobayashi H, Walker GC. 2008. Molecular determinants of a symbiotic chronic infection. Annu Rev Genet 42:413–441.

7. Pierre O, Engler G, Hopkins J, Brau F, Boncompagni E, Hérouart D. 2013. Peribacteroid space acidification: a marker of mature bacteroid functioning in Medicago truncatula nodules. Plant Cell Environ 36:2059–2070.

8. Hunt S. 1993. Gas Exchange of Legume Nodules and the Regulation of Nitrogenase Activity. Annu Rev Plant Physiol Plant Mol Biol 44:483–511.

9. Tookmanian EM, Belin BJ, Sáenz JP, Newman DK. 2021. The role of hopanoids in fortifying rhizobia against a changing climate. Environ Microbiol 23:2906–2918.

10. Reckling M, Hecker J-M, Bergkvist G, Watson CA, Zander P, Schläfke N, Stoddard FL, Eory V, Topp CFE, Maire J, Bachinger J. 2016. A cropping system assessment framework—Evaluating effects of introducing legumes into crop rotations. European Journal of Agronomy 76:186–197.

11. Bullock DG. 1992. Crop rotation. CRC Crit Rev Plant Sci 11:309–326.

12. Loureiro M de F, Kaschuk G, Alberton O, Hungria M. 2007. Soybean [Glycine max (L.) Merrill] rhizobial diversity in Brazilian oxisols under various soil, cropping, and inoculation managements. Biol Fertil Soils 43:665–674.

13. Zhang NN, Sun YM, Li L, Wang ET, Chen WX, Yuan HL. 2010. Effects of intercropping and Rhizobium inoculation on yield and rhizosphere bacterial community of faba bean (Vicia faba L.). Biol Fertil Soils 46:625–639.

14. Roughley RJ, Gemell LG, Thompson JA, Brockwell J. 1993. The number of Bradyrhizobium SP. (Lupinus) applied to seed and its effect on rhizosphere colonization, nodulation and yield of lupin. Soil Biol Biochem 25:1453–1458.

15. Corich V, Giacomini A, Vendramin E, Vian P, Carlot M, Concheri G, Polone E, Casella S, Nuti MP, Squartini A. 2007. Long term evaluation of field-released genetically modified rhizobia. Environ Biosafety Res 6:167–181.

16. O’Callaghan M. 2016. Microbial inoculation of seed for improved crop performance: issues and opportunities. Appl Microbiol Biotechnol 100:5729–5746.

17. Karmakar K, Rana A, Rajwar A, Sahgal M, Johri BN. 2015. Legume-Rhizobia Symbiosis Under Stress, p. 241–258. In Arora, NK (ed.), Plant microbes symbiosis: applied facets. Springer India, New Delhi.

18. Ilangumaran G, Smith DL. 2017. Plant growth promoting rhizobacteria in amelioration of salinity stress: A systems biology perspective. Front Plant Sci 8:1768.

19. Zahran HH. 1999. Rhizobium-legume symbiosis and nitrogen fixation under severe conditions and in an arid climate. Microbiol Mol Biol Rev 63:968–89, table of contents.

20. Delgado-Baquerizo M, Oliverio AM, Brewer TE, Benavent-González A, Eldridge DJ, Bardgett RD, Maestre FT, Singh BK, Fierer N. 2018. A global atlas of the dominant bacteria found in soil. Science 359:320–325.

21. Choma A, Komaniecka I, Zebracki K. 2017. Structure, biosynthesis and function of unusual lipids A from nodule-inducing and N2-fixing bacteria. Biochim Biophys Acta Mol Cell Biol Lipids 1862:196–209.

22. Serrato RV. 2014. Lipopolysaccharides in diazotrophic bacteria. Front Cell Infect Microbiol 4:119.

23. Nikaido H. 2003. Molecular Basis of Bacterial Outer Membrane Permeability Revisited. Microbiol Mol Biol Rev 67:593–656.

24. Komaniecka I, Zamłyńska K, Zan R, Staszczak M, Pawelec J, Seta I, Choma A. 2016. Rhizobium strains differ considerably in outer membrane permeability and polymyxin B resistance. Acta Biochim Pol 63:517–525.

25. Kulkarni G, Busset N, Molinaro A, Gargani D, Chaintreuil C, Silipo A, Giraud E, Newman DK. 2015. Specific hopanoid classes differentially affect free-living and symbiotic states of Bradyrhizobium diazoefficiens. MBio 6:e01251–15.

26. Belin BJ, Busset N, Giraud E, Molinaro A, Silipo A, Newman DK. 2018. Hopanoid lipids: from membranes to plant-bacteria interactions. Nat Rev Microbiol 16:304–315.

27. Silipo A, Vitiello G, Gully D, Sturiale L, Chaintreuil C, Fardoux J, Gargani D, Lee H-I, Kulkarni G, Busset N, Marchetti R, Palmigiano A, Moll H, Engel R, Lanzetta R, Paduano L, Parrilli M, Chang W-S, Holst O, Newman DK, Garozzo D, D’Errico G, Giraud E, Molinaro A. 2014. Covalently linked hopanoid-lipid A improves outer-membrane resistance of a Bradyrhizobium symbiont of legumes. Nat Commun 5:5106.

28. Busset N, Di Lorenzo F, Palmigiano A, Sturiale L, Gressent F, Fardoux J, Gully D, Chaintreuil C, Molinaro A, Silipo A, Giraud E. 2017. The Very Long Chain Fatty Acid (C26:25OH) Linked to the Lipid A Is Important for the Fitness of the PhotosyntheticBradyrhizobiumStrain ORS278 and the Establishment of a Successful Symbiosis withAeschynomeneLegumes. Front Microbiol 8:1821.

29. Komaniecka I, Choma A, Mazur A, Duda KA, Lindner B, Schwudke D, Holst O. 2014. Occurrence of an unusual hopanoid-containing lipid A among lipopolysaccharides from Bradyrhizobium species. J Biol Chem 289:35644–35655.

30. Belin BJ, Tookmanian EM, de Anda J, Wong GCL, Newman DK. 2019. Extended Hopanoid Loss Reduces Bacterial Motility and Surface Attachment and Leads to Heterogeneity in Root Nodule Growth Kinetics in a Bradyrhizobium-Aeschynomene Symbiosis. Mol Plant Microbe Interact 32:1415–1428.

31. Miller JH. 1972. Experiments in Molecular Genetics. Cold Spring Harbor Laboratory Pr, Cold Spring Harbor, N.Y.].

32. Mesa S, Hauser F, Friberg M, Malaguti E, Fischer H-M, Hennecke H. 2008. Comprehensive assessment of the regulons controlled by the FixLJ-FixK2-FixK1 cascade in Bradyrhizobium japonicum. J Bacteriol 190:6568–6579.

33. Regensburger B, Hennecke H. 1983. RNA polymerase from Rhizobium japonicum. Arch Microbiol 135:103–109.

34. Sadowsky MJ, Tully RE, Cregan PB, Keyser HH. 1987. Genetic Diversity in Bradyrhizobium japonicum Serogroup 123 and Its Relation to Genotype-Specific Nodulation of Soybean. Appl Environ Microbiol 53:2624–2630.

35. Cole MA, Elkan GH. 1973. Transmissible resistance to penicillin G, neomycin, and chloramphenicol in Rhizobium japonicum. Antimicrob Agents Chemother 4:248–253.

36. Kaczmarczyk A, Vorholt JA, Francez-Charlot A. 2013. Cumate-inducible gene expression system for sphingomonads and other Alphaproteobacteria. Appl Environ Microbiol 79:6795–6802.

37. Eaton RW. 1997. p-Cymene catabolic pathway in Pseudomonas putida F1: cloning and characterization of DNA encoding conversion of p-cymene to p-cumate. J Bacteriol 179:3171–3180.

38. Eaton RW. 1996. p-Cumate catabolic pathway in Pseudomonas putida Fl: cloning and characterization of DNA carrying the cmt operon. J Bacteriol 178:1351–1362.

39. Ledermann R. 2017. Role of general stress response in trehalose biosynthesis for functional rhizobia-legume symbiosis. Doctoral dissertation, ETH Zurich.

40. Gibson DG, Young L, Chuang R-Y, Venter JC, Hutchison CA, Smith HO. 2009. Enzymatic assembly of DNA molecules up to several hundred kilobases. Nat Methods 6:343–345.

41. Beck C, Marty R, Kläusli S, Hennecke H, Göttfert M. 1997. Dissection of the transcription machinery for housekeeping genes of Bradyrhizobium japonicum. J Bacteriol 179:364–369.

42. Masloboeva N, Reutimann L, Stiefel P, Follador R, Leimer N, Hennecke H, Mesa S, Fischer H-M. 2012. Reactive oxygen species-inducible ECF σ factors of Bradyrhizobium japonicum. PLoS One 7:e43421.

43. Hahn M, Meyer L, Studer D, Regensburger B, Hennecke H. 1984. Insertion and deletion mutations within the nif region of Rhizobium japonicum. Plant Mol Biol 3:159–168.

44. Ledermann R, Bartsch I, Remus-Emsermann MN, Vorholt JA, Fischer H-M. 2015. Stable Fluorescent and Enzymatic Tagging of Bradyrhizobium diazoefficiens to Analyze Host-Plant Infection and Colonization. Mol Plant Microbe Interact 28:959–967.

45. Tjørve KMC, Tjørve E. 2017. The use of Gompertz models in growth analyses, and new Gompertz-model approach: An addition to the Unified-Richards family. PLoS One 12:e0178691.

46. Zwietering MH, Jongenburger I, Rombouts FM, van’t Riet K. 1990. Modeling of the bacterial growth curve. Appl Environ Microbiol 56:1875–1881.

47. Levental KR, Malmberg E, Symons JL, Fan Y-Y, Chapkin RS, Ernst R, Levental I. 2020. Lipidomic and biophysical homeostasis of mammalian membranes counteracts dietary lipid perturbations to maintain cellular fitness. Nat Commun 11:1339.

48. Rizk S, Henke P, Santana-Molina C, Martens G, Gnädig M, Nguyen NA, Devos DP, Neumann-Schaal M, Saenz JP. 2021. Functional diversity of isoprenoid lipids in Methylobacterium extorquens PA1. Mol Microbiol.

49. Clifton LA, Skoda MWA, Le Brun AP, Ciesielski F, Kuzmenko I, Holt SA, Lakey JH. 2015. Effect of divalent cation removal on the structure of gram-negative bacterial outer membrane models. Langmuir 31:404–412.

50. Schäfer A, Tauch A, Jäger W, Kalinowski J, Thierbach G, Pühler A. 1994. Small mobilizable multi-purpose cloning vectors derived from the Escherichia coli plasmids pK18 and pK19: selection of defined deletions in the chromosome of Corynebacterium glutamicum. Gene 145:69–73.

51. Bolan NS, Hedley MJ, White RE. 1991. Processes of soil acidification during nitrogen cycling with emphasis on legume based pastures. Plant Soil 134:53–63.

52. Sáenz JP, Sezgin E, Schwille P, Simons K. 2012. Functional convergence of hopanoids and sterols in membrane ordering. Proc Natl Acad Sci USA 109:14236–14240.

53. Ma Y, Benda A, Kwiatek J, Owen DM, Gaus K. 2018. Time-Resolved Laurdan Fluorescence Reveals Insights into Membrane Viscosity and Hydration Levels. Biophys J 115:1498–1508.

54. Steinkühler J, Sezgin E, Urbančič I, Eggeling C, Dimova R. 2019. Mechanical properties of plasma membrane vesicles correlate with lipid order, viscosity and cell density. Commun Biol 2:337.

55. Zgurskaya HI, Löpez CA, Gnanakaran S. 2015. Permeability Barrier of Gram-Negative Cell Envelopes and Approaches To Bypass It. ACS Infect Dis 1:512–522.

56. Sáenz JP, Grosser D, Bradley AS, Lagny TJ, Lavrynenko O, Broda M, Simons K. 2015. Hopanoids as functional analogues of cholesterol in bacterial membranes. Proc Natl Acad Sci USA 112:11971–11976.

57. Brandenburg K, Seydel U. 1990. Investigation into the fluidity of lipopolysaccharide and free lipid A membrane systems by Fourier-transform infrared spectroscopy and differential scanning calorimetry. Eur J Biochem 191:229–236.

58. Welander PV, Hunter RC, Zhang L, Sessions AL, Summons RE, Newman DK. 2009. Hopanoids play a role in membrane integrity and pH homeostasis in Rhodopseudomonas palustris TIE-1. J Bacteriol 191:6145–6156.

59. Vitiello G, Oliva R, Petraccone L, Vecchio PD, Heenan RK, Molinaro A, Silipo A, D’Errico G, Paduano L. 2021. Covalently bonded hopanoid-Lipid A from Bradyrhizobium: The role of unusual molecular structure and calcium ions in regulating the lipid bilayers organization. J Colloid Interface Sci 594:891–901.

60. Papahadjopoulos D, Portis A, Pangborn W. 1978. Calcium-induced lipid phase transitions and membrane fusion. Ann N Y Acad Sci 308:50–66.

61. Macció D, Fabra A, Castro S. 2002. Acidity and calcium interaction affect the growth of *Bradyrhizobium* sp. and the attachment to peanut roots. Soil Biol Biochem 34:201–208.

62. Izmailov SF. 2003. Calcium-Based Interactions of Symbiotic Partners in Legumes: Role of Peribacteroid Membrane. Russian Journal of Plant Physiology.

63. Liu J, Miller SS, Graham M, Bucciarelli B, Catalano CM, Sherrier DJ, Samac DA, Ivashuta S, Fedorova M, Matsumoto P, Gantt JS, Vance CP. 2006. Recruitment of novel calcium-binding proteins for root nodule symbiosis in Medicago truncatula. Plant Physiol 141:167–177.

64. Andreev IM, Andreeva IN, Dubrovo PN, Krylova VV, Kozharinova GM, Izmailov SF. 2001. Calcium Status of Yellow Lupin Symbiosomes as a Potential Regulator of Their Nitrogenase Activity: The Role of the Peribacteroid Membrane. Russian Journal of Plant Physiology.

65. Los DA, Murata N. 2004. Membrane fluidity and its roles in the perception of environmental signals. Biochim Biophys Acta 1666:142–157.

66. Miller KJ, Wood JM. 1996. Osmoadaptation by rhizosphere bacteria. Annu Rev Microbiol 50:101–136.

67. Tejima K, Arima Y, Yokoyama T, Sekimoto H. 2003. Composition of amino acids, organic acids, and sugars in the peribacteroid space of soybean root nodules. Soil Sci Plant Nutr 49:239–247.

68. Mangiarotti A, Genovese DM, Naumann CA, Monti MR, Wilke N. 2019. Hopanoids, like sterols, modulate dynamics, compaction, phase segregation and permeability of membranes. Biochim Biophys Acta Biomembr 1861:183060.

69. Kannenberg E, Blume A, McElhaney RN, Poralla K. 1983. Monolayer and calorimetric studies of phosphatidylcholines containing branched-chain fatty acids and of their interactions with cholesterol and with a bacterial hopanoid in model membranes. Biochimica et Biophysica Acta (BBA) - Biomembranes 733:111–116.

70. Chen Z, Sato Y, Nakazawa I, Suzuki Y. 1995. Interactions between bacteriohopane-32,33,34,35-tetrol and liposomal membranes composed of dipalmitoylphosphatidylcholine. Biol Pharm Bull 18:477–480.

71. Crowe LM, Mouradian R, Crowe JH, Jackson SA, Womersley C. 1984. Effects of carbohydrates on membrane stability at low water activities. Biochim Biophys Acta 769:141–150.

72. Casadaban MJ, Cohen SN. 1980. Analysis of gene control signals by DNA fusion and cloning in Escherichia coli. J Mol Biol 138:179–207.

73. Simon R, Priefer U, Pühler A. 1983. A Broad Host Range Mobilization System for In Vivo Genetic Engineering: Transposon Mutagenesis in Gram Negative Bacteria. Nat Biotechnol 1:784–791.

74. Ledermann R, Strebel S, Kampik C, Fischer H-M. 2016. Versatile Vectors for Efficient Mutagenesis of Bradyrhizobium diazoefficiens and Other Alphaproteobacteria. Appl Environ Microbiol 82:2791–2799.

